# Threat-Related Corticocortical Connectivity Elicited by Rapid Auditory Looms

**DOI:** 10.1101/2024.06.21.600106

**Authors:** Karolina Ignatiadis, Roberto Barumerli, Gustavo Deco, Brigitta Tóth, Robert Baumgartner

## Abstract

While sounds of approaching objects are generally more salient than those of receding ones, the traditional association of this auditory looming bias with threat perception is subject to debate. Differences between looming and receding sounds may also be learned through non-threatening multisensory information, or influenced by confounding stimulus characteristics. To investigate, we analyzed corticocortical connectivity patterns from electroencephalography, examining the preferential processing of looming sounds under different attentional states. To simulate rapid distance changes we used complementary distance cues, previously studied in the looming bias literature. Notably, despite the absence of conscious threat perception, we observed crucial involvement of frontal cortical regions typically associated with threat and fear responses. Our findings suggest an underlying bias towards the ventral ‘what’ stream over the dorsal ‘where’ stream in auditory information processing, even when the participants’ task was solely focused on the discrimination of movement direction. These results support the idea, that the perceptual bias towards looming sounds reflects an auditory threat detection mechanism, while offering insights into the neural function involved in processing ecologically relevant environmental cues.

## 1 Introduction

If a car is approaching from a distance, its timely detection and avoidance are essential to our survival. It is presumably to improve warning capacity, that approaching stimuli are more salient than receding ones. This perceptual asymmetry, often referred to as the auditory looming bias, has been found present across species [1–5] and ages [6–9], making it a rather universal trait. As an effective warning mechanism, the looming bias should have the capacity to readily capture attention and be rather universal across cue types. Corroborating this hypothesis, signatures of the bias have indeed been found across attentional states and auditory distance cue types; they were present already at the level of Heschl’s gyrus (HG), housing the primary auditory cortex [10]. Yet the bias’ relationship to threat detection remained a hypothesis, and discrepancies in behavioral performance, timing, and attentional amplification, suggest that there are differences in cortical processing depending on these factors.

The notion of stimulus-specificity has frequently been put forth with regards to the selective advantage of the looming bias, and its function as a warning mechanism for organisms facing potential collisions with sound sources. It tends to be observed more consistently in response to stimuli with a natural overtone structure, in contrast to Gaussian white noise stimuli, which sound arguably more artificial [4, 9, 11]. However, studies have also demonstrated looming biases in response to noise stimuli when accounting for the natural acoustic filtering properties of listeners [12, 13]. This suggests that the absence of natural spatial cues, rather than the source’s identity, may be responsible for the failure to elicit the bias in certain cases. Additionally, some investigations into the looming bias have employed auditory distance changes as short as 10 ms [10, 12, 13], prompting questions about the necessary identity and ecological validity required to evoke this effect.

From a neuroimaging perspective, increased amygdala activation in response to slowly rising sound intensities has been an important argument for the bias’ warning function [14]. Apart from Heschl’s gyrus and the amygdala, functional magnetic resonance imaging (fMRI) has highlighted the involvement of the temporal plane, superior temporal sulcus (STS), prefrontal cortex (PFC), and inferior parietal lobe (IPL) in the preferential processing of looming sounds [14, 15]. In general, auditory stimuli have been hypothesized to follow two parallel cortical processing streams: one following a ventral and the other a dorsal path [16]. Originally stemming from visual research, the dorsal pathway is associated with spatial perception (”where”), while the ventral stream with object identification (”what”) [16, 17]. The inferior parietal lobe, as part of the dorsal auditory pathway, is thought to play a crucial role in spatial hearing [18, 19] and sound motion processing in particular [20], whereas superior temporal sulcus and prefrontal cortex belong to the ventral pathway. Based on these findings, the looming bias circuit emerges as an extended distributed cortical network.

Besides the mere activation increase induced by looming sounds, one crucial aspect is the way in which the involved regions are at interplay. This question can be addressed through functional connectivity investigations [21]; namely computational methods exploring the information exchange among regions of interest (ROIs). Unlike structural connectivity, which describes anatomical connections linking sets of neural elements, functional connectivity is dynamic in nature. It represents changes in statistical inter-dependencies between or among brain regions, within a specific time interval and connected to an event of interest. The observed brain regions that are found to contribute the most with regards to connectivity, relative to the others or the combinations thereof, are defined as functional hubs. Those are also dynamic and may deviate from an anatomical definition, as they can be a part of different functional clusters [22]. Findings stemming from available connectivity analyses of the auditory looming bias circuit are inconclusive: a study based on intensity ramps argues that top-down directional causal influence from prefrontal cortex to Heschl’s gyrus enhances processing of looming versus receding sounds [23], while prior investigations on spectral stimuli argue for a bottom-up, temporofrontal connectivity [13].

Functional connectivity methods employed in previous research focus on a bidirectional analysis process, albeit relying on a small preselection of brain regions (Granger [24], Phase Transfer Entropy [25]). Although insightful regarding the interplay of the considered ROI pairs, further methods may offer an approach that is closer to a network structure. They are nevertheless limited by either the number of regions that can be considered (conditional Granger Causality [26]), the dimensionality (multivariate Granger [27]) or the amount of constraints enforced by parameters of a model (e.g., Dynamic Causal Modeling [28]). Contrary to that, recent frameworks offer both a holistic as well as data-driven approach [29, 30]. They provide the possibility to investigate the whole brain on different levels, without the necessity of a predefined set of ROIs or network structure parameters. Of those, the INSIDEOUT approach [29] relies on the observation, that the environment drives hierarchically lower, sensory regions, stronger than hierarchically higher ones. A system in equilibrium has seamless transitions between different states; thermodynamically, it is reversible in time. Should the system get driven out of equilibrium, the transitions between states become non-reversible and an arrow of time emerges. Measuring the effects of the extrinsic environment on the intrinsic brain dynamics through the non-reversibility of the system, here the brain, can therefore help uncover variations in brain states under different conditions. The framework of normalized directed transfer entropy (NDTE) [30], contrarily, works on a mesoscopic level: By considering the interconnectivity of all defined brain regions, it draws assumptions about the most essential contributors, or functional hubs, of the underlying networks.

In the current study, we investigate the cortical connectivity network underlying the auditory looming bias under the individual factors of cue type and attention, in search for overlapping patterns along spatial and/or identity-related cortical processing streams. Through the high temporal resolution of electroencephalography (EEG) in combination with recently proposed, data-driven approaches for connectivity analyses, we investigate the brain at different levels of granularity [29, 30]: First as a whole, and subsequently in search for the functional hubs that act as essential contributors in the looming network. High spatial resolution is achieved by complementing source localisation of high-density EEG with individual brain anatomies and electrode locations [31]. The present analyses are based on prior collected data, studying the auditory looming bias at the level of HG under the aforementioned factors of attention and cue type [10]. In that paradigm (Fig. 1), participants listened to broadband harmonic tones that rapidly changed in their simulated distance from the listener and thereby elicited a looming or receding percept. Distance cues comprised either overall sound intensity or spectral shape changes. Listeners were first passively exposed to the stimuli while watching a silent subtitled movie and later had to discriminate the sonic motion direction. We find that there is to be higher sensitivity for intensity stimuli, while different main hubs, traditionally connected to threat and fear perception, emerge based on the factors considered.

**Fig. 1:**
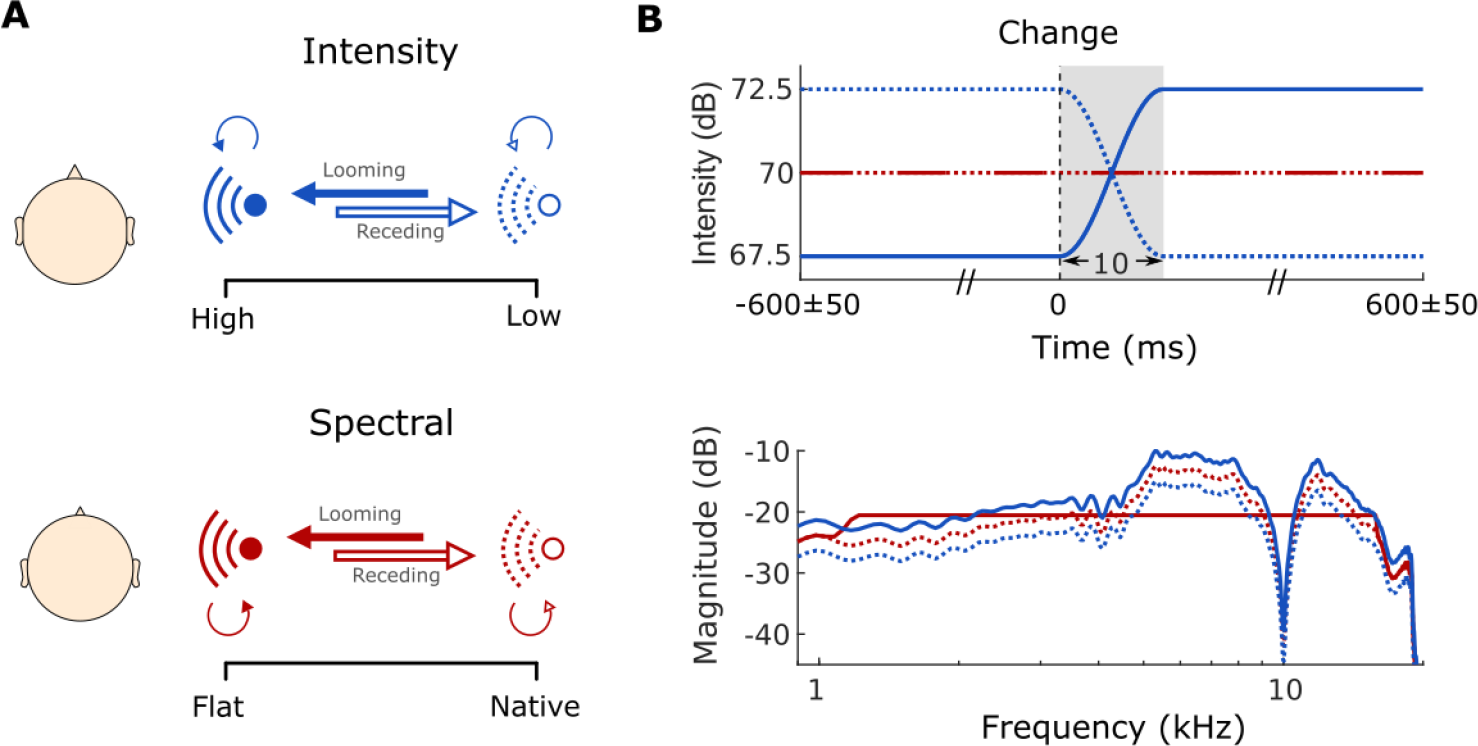
Experimental design. **A**) Looming and receding percepts created through simulated transition between two sounds of different intensities (top, blue) or spectral shapes (bottom, red). Thick arrows represent 50% transition probability for motion trials (dark = looming; light = receding). **B**) Sound intensity over time (top panel) and magnitude spectrum (bottom panel) of all implemented stimuli. Figure adapted from [10].

## 2 Results

For the present connectivity investigations we extracted the source-localized EEG time series of all cortical regions, as defined by the Desikan-Killiany parcellation [32]. We considered the time interval between 0 and 300 ms relative to the event of distance change. This choice was made based on the finding, that this time window has shown significant biases evoked in HG in previous investigations [10].

### 2.1 Intensity looms induce stronger non-reversibility in cortical processing

INSIDEOUT reflects how the environment (extrinsic, outside) affects the dynamics and equilibrium of the underlying brain state (intrinsic, inside) [29], by measuring the non-reversibility of a considered system.

We implemented this framework by accounting for the set of all ROIs of the considered parcellation, hence the cortex as a whole (Sec. 4.4.1). Higher non-reversibility is thus understood as a quantification of the amount of change in causal interactions of the brain under each considered condition.

Figure 2 shows the distributions of non-reversibility measures obtained for every considered condition. An ANOVA with the factors of attention, cue type and motion direction revealed the latter to be a significant factor (*F*_1,27_ = 8.34, *η*^2^ = 0.24, *p* = 0.008), as well as its interaction with the cue type (*F*_1,27_ = 5.11, *η*^2^ = 0.16, *p* = 0.032). To further investigate this interaction, we performed a separate ANOVA for each cue type and adjusted the p-values for multiple comparisons using Bonferroni correction. For intensity, looming sounds were found to elicit higher non-reversibility than receding sounds (*F*_1,27_ = 9.17, *η*^2^ = 0.25, *p* = 0.011). No significant factors or interactions thereof appeared for the spectral condition. Hence, looming stimuli appeared to disrupt the intrinsic equilibrium more than receding ones, in particular when they are based on intensity changes.

**Fig. 2:**
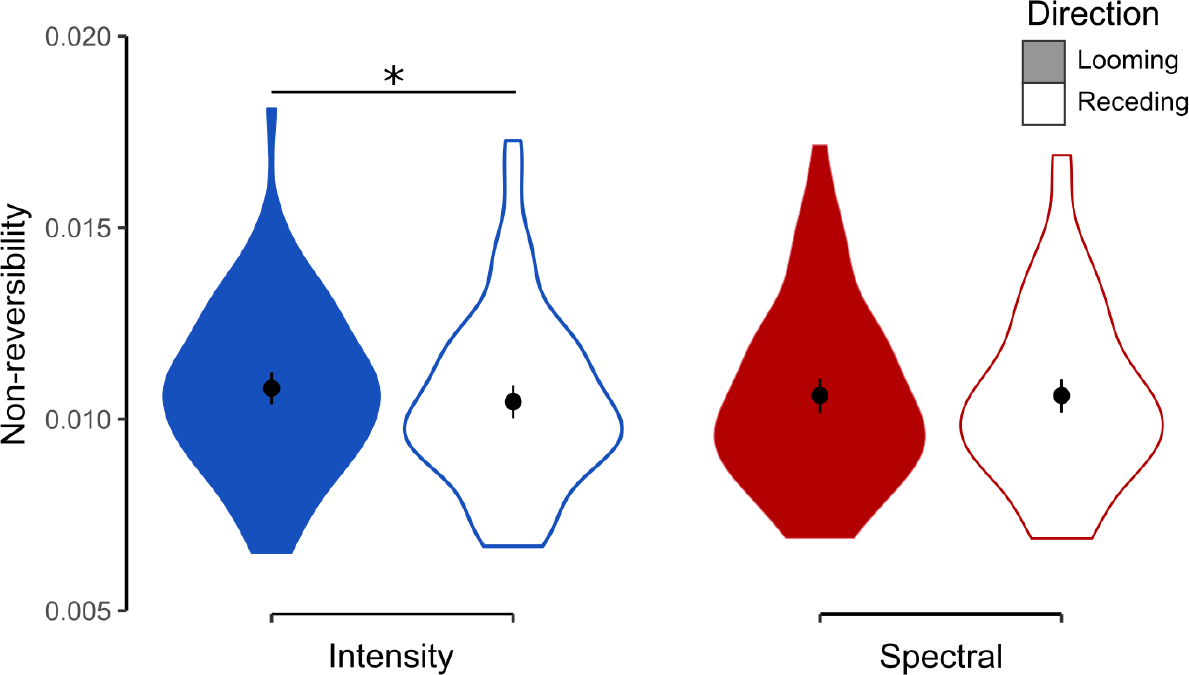
Effects of cue type and movement direction on the temporal non-reversibility of cortical processing, as calculated by the INSIDEOUT framework. Points and bars represent the means and their standard errors within each violin plot. Asterisks indicate statistical significance (p < 0.05).

### 2.2 Connectivity hubs relevant to the auditory looming bias

To better understand the dependencies, we then applied the NDTE framework, as it offers a more granular view on the interacting brain regions. Following the procedure suggested in [30], we considered each ROI’s connection to every other ROI in the cortical parcellation. The connectivity between each pair of regions was calculated on the actual data, and its significance assessed through a distribution of surrogate data stemming from the same ROI-pair. Aggregation of the connectivity information across subjects allowed for the construction of connectivity hubs, namely regions, or sets thereof, that are, as a whole, more connected compared to any other considered set comprising the same number of ROIs. We performed a connectivity analysis on the bias data by considering the factors of attention and cue type.

The two quantities of essence in this framework are termed inflow (*G_in_* in [30]) and outflow (*G_out_* in [30]); they respectively represent the connectivity incoming to or outflowing from a ROI. If a set of ROIs is considered as a network, inflow is the sum of all incoming connectivity across all its constituent ROIs. The respective holds for the outflow.

We determined the major inflow and outflow hubs per condition by following the concept and search procedure of functional rich clubs (Sec. 4.4.2; FRICs in [30]). Following the procedure for their definition based on the inflow, we respectively defined the major hubs based on the connectivity outflow (Fig. 3, Sec. 4.4.2). As demonstrated in figure 3A, the major inflow was attributed to one region, except for the active intensity condition. The ROIs receiving the most inflow spanned over temporal regions (STG), frontal regions (pars opercularis - IFGoper, frontal pole - Fpole), and both hemispheres. Across both active conditions, only the frontal pole emerged as a crucial inflow hub for looming bias. In the passive conditions, the relevant inflow hubs comprised the right precentral gyrus (PreCG) for intensity and the left pars triangularis (IFGtriang) for spectral stimuli. Regarding the outflow hubs, one region emerged per condition and all regions were located in the left hemisphere. Apart from the active intensity condition, where the insular gyrus (IG) was identified as the main hub, temporal regions were identified for the remaining cases: superior temporal gyrus (STG) for active spectral, transverse temporal gyrus (Heschl’s Gyrus, HG) for passive intensity, and the banks of the superior temporal sulcus (BanksSTS) for passive spectral.

**Fig. 3:**
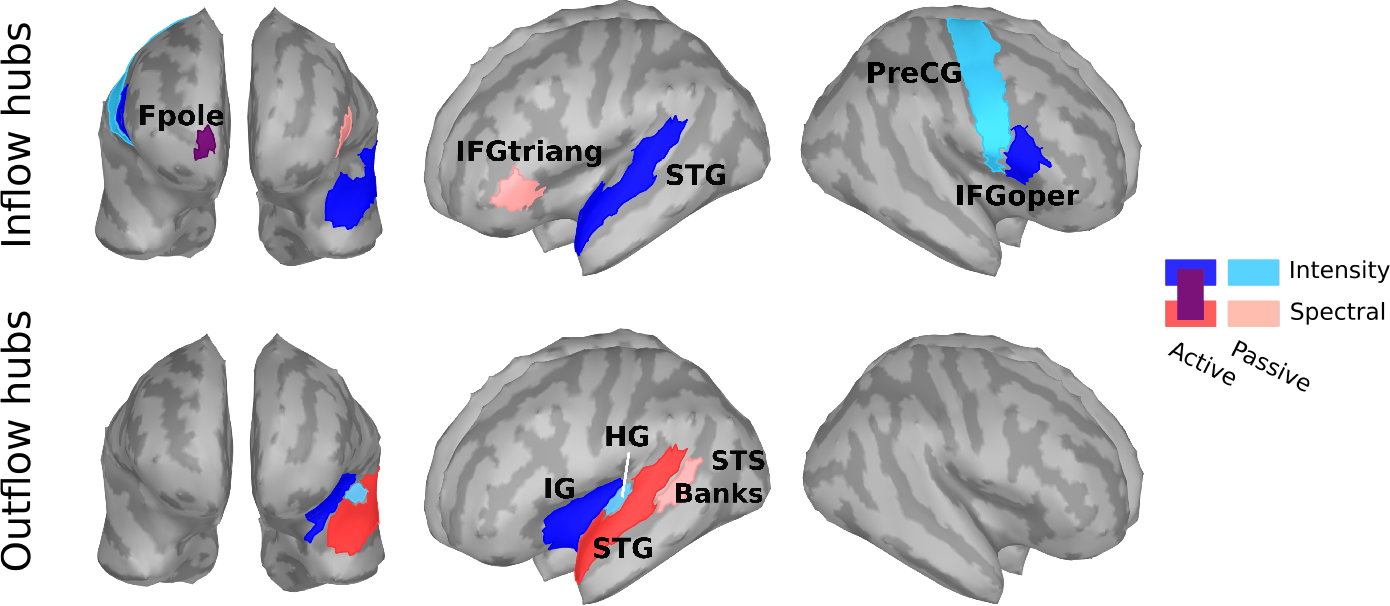
Major inflow and outflow hubs of looming bias identified per considered condition. Fpole (magenta) is activated by both active intensity and spectral. Fpole - frontal pole, IFGtriang - pars triangularis, STG - superior temporal gyrus, PreCG - precentral gyrus, IFGoper - pars opercularis, IG - insular gyrus, BanksSTS - banks of the superior temporal sulcus, HG - Heschl’s gyrus.

We further extracted the pattern of hub connections that emerged, separately for inflow and outflow connectivity in each considered condition (Fig. 4). On a large scale, the inflow hubs, localized to the frontal cortices (Fpole, IFG, PreCG), dominantly received information from more distant regions of the sensory temporal regions such as the superior temporal gyrus, Heschl’s gyrus or inferior temporal cortices. In contrast, the outflow hubs, localized to the temporal regions (IG, STG, HG, STS), tended to send information to more local areas within the temporal cortex. All those connections occurred mainly within hemisphere, reflecting rather weak inter-hemispheric connections to and from the hubs.

**Fig. 4:**
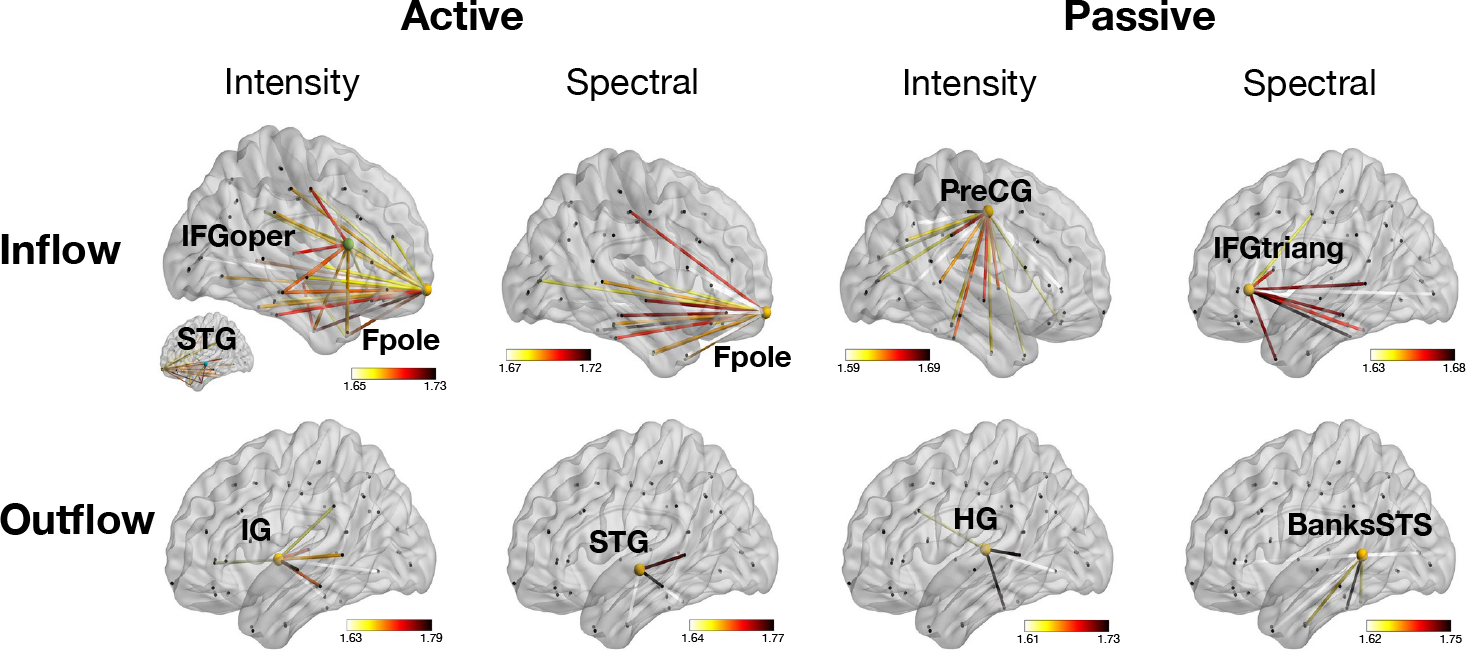
Detailed connectivity for major inflow and outflow hubs identified per condition. Displayed connections correspond to the top 30% of connectivity strength per condition. Color bars were adjusted accordingly and normalized connectivity values were scaled up by a factor of 100.

## 3 Discussion

Looming sounds exhibit remarkable salience, consistently eliciting a perceptual bias compared to equivalent receding sounds. This bias is commonly hypothesized to signify a mechanism for threat detection and hazard protection. In this study, we employed two data-driven, state-of-the-art functional connectivity approaches, to examine cortical responses to simulations of rapid changes in auditory distance. We examined the responses during both passive and active listening, and while utilizing either intensity or spectral cues. Analyzing macroscopic brain states, temporal non-reversibility tests revealed a more pronounced impact of looming sounds on the overall functional connectivity when intensity was employed to simulate motion in distance. Taking a more granular approach, we identified functional connectivity hubs for each condition, shedding light on the intricate neural networks underlying these perceptual processes. Throughout our conditions, frontal regions emerged as the main inflow hubs (frontal pole, IFG including the pars opercularis, pars triangularis also known as Broca’s area-BA45), and temporal regions as the major outflow hubs (primary and secondary auditory cortices such as the superior temporal gyrus, transverse temporal gyrus including the PAC, banks STS). Prior studies have argued regarding the directionality of the connectivity in the looming bias, with results divided between bottom-up processing [13] or a top-down intervention [23]. The here-identified hubs support the bottom-up notion, as temporal regions act more as sources (outflow hubs) and frontal regions as receivers (inflow hubs) in response to looming sounds. Moreover, those hubs mostly appear to belong to the ventral auditory pathway. As seen in the case of passive intensity, the motor cortex is additionally present, while no hubs appear in our findings that could be clearly associated with the dorsal auditory pathway. This result is intriguing, as it suggests that, in the considered context, priority is given to recognizing and identifying the auditory sound source (”what”) rather than its location (”where”); even though listeners were solving a spatial auditory task during the active listening part of their study participation. The to be expected high involvement of the dorsal pathway in spatial perception might have been canceled out by the bias calculation (difference between looming and receding). The importance of source identity on the elicitation of auditory looming bias has previously also been demonstrated by comparisons between different types of source stimuli: tones elicited stronger biases than noise [4, 5, 9, 33, 34]. The here identified connectivity hubs seem to reflect the crucial role of source identity in the manifestation of the auditory looming bias.

### 3.1 Emerging looming bias hubs relate to fear responses

The analysis of sound source identity is governed by cortical hubs that are congruent to various literature findings in the context of threat and fear. Among the frontal regions, the anterior part of the frontal cortex (Fpole) emerged as the major inflow hub in both active conditions, which is in agreement with its recognized function for executive cognitive processing and action selection [35, 36]. Other auditory studies also demonstrated its involvement in the context of threatening sentences and emotionally salient pictures [37]. Apart from the frontal pole, parts of the inferior frontal gyrus appear as essential in facilitating the bias: the right pars orbitalis for the active intensity and left pars triangularis for passive spectral. More generally, studies of visually presented threat-related words have reported activation of the left inferior frontal gyrus [37, 38]. Inferior frontal gyrus has additionally been activated in the context of fear conditioning [39] and the downregulation of psychophysiological reactions to threat [40].

The information received by the major inflow hubs primarily appeared as coming from primary and secondary auditory cortical regions on the temporal lobe. Concordantly, the main outflow hubs we identified were localized in the temporal areas. Being an auditory task, areas such as the PAC may be especially involved; yet previous research has also implicated it in threatening sound paradigms [37]. Superior temporal ROIs, emerging in our considered active (attentive) conditions, have additionally been connected to attention linked to threatening voices [41].

In contrast to mainly temporal regions as outflow hubs, the insula appeared as a key outflow node in the intensity stimuli. As a ROI, it has been associated with fear and anxiety conditioning [37, 39, 42], while animal studies have demonstrated its implication in fear or extinction memory [43]. Intensity stimuli have behaviorally and neurally emerged as more salient than their spectral counterparts, in both adult and newborn listeners [10]. Implication of the insula only in the perception of intensity stimuli might act as a contributor of that manifestation.

Finally, frontotemporal activations have generally been linked to the basolateral amygdala (BLA), an essential hub of the limbic system, in the context of automatic fear detection [44]. The amygdala itself has, in turn, been further implicated in the looming perception [14] in a warning role. As we conducted our study with the use of EEG, subcortical activations, consequently also the amygdala, are either inaccessible or unreliable; the direct verification of the BLA-frontotemporal link in the context of the bias can thus not be made through our findings. Yet the emerging frontal and temporal connectivity hubs may be a manifestation of the BLA-frontotemporal exchange, congruently to previous findings: Invasive studies on animals have specifically implicated the medial prefrontal cortex and BLA in the discrimination between harmful and safe stimuli, and highlighted that the corticocortical dialogue between sensory and prefrontal areas is essential for fear-discrimination processes [45]. Taken together, the functional relevance of the major hubs we identified along the ventral auditory pathway suggests that, regardless of cue type, looming sounds elicit the perceptual bias by rapidly recognizing the sound as a potential threat.

### 3.2 Methodology and limitations

In the current study, we utilized direct (NDTE) and indirect (INSIDEOUT) connectivity metrics in order to obtain an image of the bias-related processes on the cortical surface. Depending on the method at hand, investigations can be done at different levels of granularity.

INSIDEOUT captures the breaking of causal connections through non-reversibility and the arrow of time in order to measure brain connectivity. Compared to other approaches, it has the big advantage that no underlying constraints (e.g., ROIs or networks) or models (e.g., directionality or node assumptions) are necessary for its implementation. It can additionally give a coarse representation of the different brain states based on the whole cortex in a significantly less computationally complex and time-consuming manner than conventional approaches would demand. In terms of non-reversibility, the looming bias was found mainly for the intensity stimuli. Broadly considered in looming studies, intensity stimuli have generally appeared more salient than spectral ones; the latter seem to be more complex in their understanding and cognitively processed in a much more subtle manner [46–48]. As INSIDEOUT is reflective of subjective conscious awareness [29], our result corroborates the difference in perception depending on cue type. The greater intervention of intensity stimuli, in terms of disruption in causal interactions, highlights their salience as already emerged through prior behavioral as well as neural studies [6, 7, 9, 10, 12, 14, 15, 23, 49, 50]. The effects we found from the INSIDEOUT framework, although present, are small in size. This is likely due to our highly specific paradigm (auditory looming bias), rendering the brain states only subtly, but not fundamentally, different. Despite this highly specific approach, though, INSIDEOUT still revealed significant effects in line with previous findings.

Contrary to the coarse granularity offered by INSIDEOUT, the fine-grained method of NDTE yielded insights into which regions are the main hubs in manifesting the looming bias, and does so in a data-driven way. By considering all ROIs of a given parcellation, the cross-connectivity is calculated. By, then, ranking regions based on their outflow (sources) or inflow (receivers) and iteratively comparing networks (Sec. 4.4.2), conclusions about ROIs, or networks thereof, with the most essential contribution per considered condition emerge. It should be noted that the timescale of all effects is defined by the calculated minimum of the autocorrelation function. As shown in previous research, this is a solid approach to our investigations [29, 30]. In a more ideal way, though, and although computationally significantly more costly, this parameter could be set individually for each considered time series.

In our investigation we adhered to the rather coarse parcellation of the Desikan-Killiany atlas [32]. Our selection relies on both aiming to compare outcomes to prior literature [13, 23] as well as reduce complexity, especially in the case of NDTE calculations. Finer parcellations, such as the one from Destrieux [51], could offer different insights depending on the question at hand; yet they come with higher amount of regions and therefore complexity. Finer parcellations may additionally be more prone to wrongful activity attribution if the precision of the source localization is insufficient. Although we used individualised anatomical information to aid the performed EEG source localization [31], spatial imprecisions are inevitable. An example thereof is the depth-weighting done by algorithms for sources that are intricately placed on the cortex. Additionally, should activity arise from subcortical surfaces at greater distances from the sensors, EEG may wrongfully attribute the recorded activity. Our results are in good agreement with relevant literature, yet different imaging methods, selected parcellations or implemented algorithms may lead to slightly altered outcomes.

## 4 Methods

### 4.1 Participants

Thirty-five healthy young adults were invited for study participation. Exclusion criteria during recruitment comprised self-reported indications of psychological and neurological disorders or acute or chronic heavy respiratory diseases that may prevent the participant from sitting still during the EEG recording.

All invited participants signed informed consent prior to testing, were neither deceived nor harmed in any way and were informed that they could abort the experiment at any time without any justification or consequences. The study was conducted in accordance with the standards of the Declaration of Helsinki (2000). No additional ethics committee approval was required given the non-medical non-invasive nature of our study, as per the Austrian Universities Act of 2002. In total, experiments lasted around five hours per participant and participants received monetary compensation in return for their time.

Prior to conducting the experiment, participants’ hearing thresholds between 1 and 12.5 kHz were measured via pure tone audiometry (Sennheiser HDA200; AGRA Expsuite application [52]), with a deviation of more than 20 dB from the age mean [53] leading to subject exclusion. Six participants were excluded (29 remaining subjects, 15 females: 25.0 ± 2.60 years old (mean ± standard deviation); 14 males: 25.1 ± 2.77 years old). An error rate in recognition of static sounds (catch-trials) exceeding 20% resulted in one additional exclusion (female, 45.2% errors).

In total, 28 participants were included in this study. They were informed of the procedure and their rights (no deception nor harm, freedom to interrupt the experiment without justification or repercussions) and signed informed consent prior to testing. The study was conducted in accordance with the standards of the Declaration of Helsinki. No additional ethics committee approval was required given the non-medical non-invasive nature of our study, as per the Austrian Universities Act of 2002. Experiments lasted around five hours per subject and participants were remunerated after testing.

### 4.2 Stimuli

The auditory stimuli were complex harmonic tones [54] (*F*_0_ = 100 Hz, bandwidth 1 − 16 kHz), filtered with listener-specific head-related transfer functions (HRTFs) to sound as coming from either the right or left direction on the interaural axis when presented over earphones. Stimulus duration was 1.2 s with 10 ms onset and offset ramps of raised-cosine shape. Inter-stimulus intervals lasted 500 ms. Trials were randomized throughout the experiment and balanced over blocks, with 50% looming and 50% receding sounds. Those were created by either modifying the intensity or the spectral shape of a sound and crossfading between the final simulated sound source positions (from far to near for the looming, and near to far for the receding condition). The intensity manipulation resulted in a sound appearing to recede while its intensity decreased with time. We presented sounds crossfading between +2.5 dB (near position) and −2.5 dB (far position) to induce looming and receding sensations (e.g., 11). For changes in spectral shape, we manipulated the individually recorded HRTFs following the procedure introduced in [12]. The different cue types were applied block-wise. Apart from movement and spatial cue type, we block-wise manipulated whether the sound source was presented from the left or the right side of the listener. The experiment consisted of two parts: an initial passive listening part, during which subjects were watching a silent subtitled movie while being exposed to 600 trials and a subsequent active part, in which subjects performed a spatial discrimination task on the presented sounds.

Stimuli and experimental procedures were programmed in MATLAB (R2018b, Mathworks, Natick, Massachusetts) with the use of the Auditory Modeling Tool-box [55] and Psychtoolbox [56].

### 4.3 Recordings and processing

EEG recordings were done with a 128-channel system (actiCAP with actiCHamp; Brain Products GmbH, Gilching, Germany) at a sampling rate of 1 kHz. Noisy channels were being noted during the recordings. All saved EEG data were visually inspected to detect potential additional noisy channels, which were then spherically interpolated. Inspected data were bandpass-filtered between 0.5 − 100 Hz (Kaiser window, *β* = 7.2, *n* = 462) and epoched ([−200, 1500] ms) relative to stimulus onset. We applied hard thresholds at −200 and 800 *µV* to detect and inspect extremely noisy trials. An additional check for the identification of additional bad channels was implemented, via an automatic channel rejection step; detected channels would then be visually inspected and interpolated. No additional noisy channels were detected for any of the subjects at this step. Independent component analysis (ICA) was followed by a manual artifact inspection and rejection of oculomotor artifacts (removal of up to 3 components per subject). The cleaned data were thereafter re-referenced to their average. Within each subject, trials were equalized to match the condition with the minimum amount within the subject after trial rejection. This was achieved within each subject by pseudo-selection aiming to maintain an equal distribution across the recordings. This resulted in an average of 569 clean trials (SD = 27.7) per subject. All preprocessing steps were undertaken on the EEGLAB free software (57) in MATLAB (R2018b, Mathworks, Natick, Massachusetts).

Twenty-five (25) of 28 participants had their individual anatomical structures and electrode positions recorded. Anatomical magnetic resonance images (MRIs) were segmented via Freesurfer [58] and used to create a study protocol on Brainstorm [59]. For the remaining 3 subjects, the default anatomical models of Brainstorm were used (ICBM152 brain template); individual MRIs could not be recorded due to incompatibilities with the scanner (suspicion of metallic parts in the body). Anatomical models were created via OpenMEEG [60] with following parameters: boundary element model (BEM) surfaces had 1922 vertices per layer for scalp, outer skull and inner skull, and a skull thickness of 4 mm. The relative conductivity was set to 0.0125 for the outer skull and to 1 for the remaining layers. Manual co-registration between head model and individual electrode locations was done for each subject individually. Recorded activity was inferred to the cortical surface via dynamic statistical parametric mapping (dSPM) [61]. The noise covariance was calculated from a 200 ms pre-stimulus interval. Dipole orientations were considered constrained to the surface and source signals were reconstructed at 15000 vertices describing the pial surface. Following previous literature [13, 23], cortical mapping was done according to the Desikan-Killiany parcellation [32]. We extracted all ROI time series from the 68 areas of the atlas, as defined in Brainstorm. Based on the evoked time courses at the level of the transverse temporal gyrus, taken from [10], a time window of 300 ms post-change was defined as the time window of interest.

### 4.4 Connectivity calculations

Our NDTE connectivity analyses, being based on Granger causality, assume stationary signals as input. In order to fulfill this stationarity requirement, we tested our time courses for this property. Following the recommendations of Brainstorm [59], each time-series was subjected to both the Kwiatkowski-Phillips-Schmidt-Shin test (KPSS) for trend-stationarity and the unit root Augmented Dickey Fuller test (ADF), as implemented in MATLAB 2018b (kpsstest, adftest; Mathworks, Natick, Massachusetts). As broadband EEG signals are highly non-stationary, stationarity of all signals was restored through double differencing of the individual time-series [24].

#### 4.4.1 INSIDEOUT

The INSIDEOUT framework [29] is based on the time-shifted correlation matrices between each considered time series and its time-reversed version, thereby echoing the asymmetry in temporal processing. The arrow of time captures the interaction with the environment: a system that remains unperturbed by external factors maintains its intrinsic equilibrium and is therefore characterised by high reversibility. Higher dissimilarity of the forward and reverse time series corresponds to higher non-reversibility, and thereby higher impact of the external environment on the intrinsic dynamics.

Reversed time series were obtained by inverting the original ones in time, for each condition, subject, ROI and trial. Correlations between time series were calculated through the MATLAB function corr, for both the forward as well as the reverse time-shifted correlations. If *FS_forward_*(*T*) and *FS_reversal_*(*T*), expressed as mutual information based on the time-shifted correlations, are the matrices representing the causal dependencies of the system, here across ROIs, the non-reversibility (non-equilibrium) per condition is calculated as

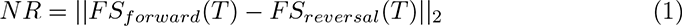

and is hence equal to the mean of the absolute squared difference between the forward and reversed matrices (cf. [29] for detailed calculations). Time-shift T is defined as the decay to the first minimum of the autocorrelation function across conditions and subjects [29, 30].

Statistical differences among the conditions were assessed based on ANOVA with the factors of attention, cue type and motion direction.

#### 4.4.2 NDTE

The data used in the assessment of NDTE were based on the extracted looming bias on a single trial basis for each ROI. All subsequent calculations were done following the pipeline described by Deco et al. [30]. In it, the statistical causal interaction between any two ROIs is assessed based on the measure of mutual information. Considering *X* and *Y* to denote the activity of the source and target ROIs, respectively, the mutual information is calculated as

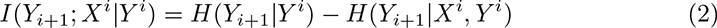

where *I*(*Y_i_*_+1_; *X^i^|Y ^i^*) corresponds to the degree of statistical dependence between the source’s past *X^i^* = [*X_i_, X_i−_*_1_, …, *X_i−_*_(_*_T−_*_1)_] and the target’s immediate future *Y_i_*_+1_ [30]. *H*(*Y_i_*_+1_*|Y ^i^*) and *H*(*Y_i_*_+1_*|X^i^, Y ^i^*) express the respective conditional entropies, that are, in the implemented framework, estimated based on covariance matrices [62]. The time interval defined by *T* stems from the autocorrelation of the time series. Following Deco et al. [30], the corresponding order (”maximum lag”) was calculated based on the decay to the first minimum of the autocorrelation function across conditions and subjects; in our case *T* = 6. In order to be able to compare and combine NDTE values across ROI pairs, calculated connectivity values were normalised as

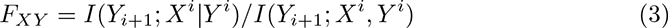

namely by the total mutual information the past of both source *X* and target *Y* hold about the future of target *Y*. This is the quantity considered throughout our calculations and yields, for each trial, an NDTE matrix with the bidirectional flow among all 68 ROIs of the parcellation.

As the whole cortical surface and all bidirectional connections therein are considered, the high amount of comparisons is susceptible to spurious correlation outcomes. For that reason, circular-shift surrogate data were generated for each considered ROI pair. P-values for each connection were assessed based on the distribution of connectivity data resulting from 100 independent circular time-shifted surrogate iterations. Statistical significance of connections between ROIs was calculated through p-value aggregation done by Stouffer’s method [63] in two steps; initially at a subject level with a within-condition aggregation across trials of a subject, and subsequently at a group level, with aggregation across all subjects of a condition. For each considered condition, the multiple comparison correction was performed by the false discovery rate method (FDR) [64]. The corrected values were then used as a binary “significance mask”, to select the significant connections per condition. The resulting data comprised one NDTE matrix of dimensions ROI x ROI per condition, containing the averaged connectivity values for the ROI pairs that survived the significance evaluations. Inflow ROIs are positioned along the first, while outflow ROIs along the second dimension of the NDTE matrix, termed *C_All_*.

For each ROI *i* of *C_All_*, the total inflow from all remaining ROIs *j* of the cortical parcellation is defined as the sum of connectivity across all columns of the matrix: *G_in_*(*i*) = Σ*_j_C_Alli,j_*. The respective holds for the total outflow per ROI *j*: *G_out_*(*j*) = Σ*_i_C_Alli,j_*.

The major hubs are identified through an iterative process. After sorting the regions based on their inflow (for inflow hubs) or outflow (for outflow hubs), an algorithm searches for the largest subset of ROIs *k* that have a value *G_hub_* significantly larger than any other set, comprising the same amount of regions. The significance value of each *G_hub_*(*k*) is assessed via 1000 Monte Carlo simulations, where for each permutation, one member of the current subset *k* is substituted with any of the remaining ones from the parcellation, and the *G_hub_*(*k*) is calculated anew. The in- and outflow values are calculated as

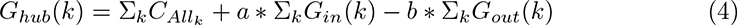

where Σ*_k_C_Allk_* is the total flow within the considered subset, Σ*_k_G_in_*(*k*) represents the total inflow to the considered subset from all ROIs of the parcellation and Σ*_k_G_out_*(*k*) the total outflow of the subset to the rest of the ROIs. For the inflow hubs the mutlipliers are [*a* = 1; *b* = 1] and for the outflow hubs [*a* = −1; *b* = −1].

By progressively adding one ROI (”node” in [30]) to the considered subset (*k* = [*i*_1_, …, *i_l_*], where *l* is the whole set of ROIs), the major hubs emerge as the set for which the in- or outflow is still within significance limits (i.e., smaller than 0.05).

## Data availability

Data are available under https://osf.io/4gdy2/.

## Code availability

Analysis scripts are available and will be updated under https://osf.io/4gdy2/. Codes for the NDTE framework [30] are available under https://github.com/decolab/nhb-ndte.

## Acknowledgments

We would like to thank Dr. Guenther Koliander for his insightful discussions on the statistical basis and background of the studied approaches, and Elvira Del Agua Banyeres for her support in the computational implementations. This research was funded by the Austrian Science Fund (FWF, I 4294-B and ZK66), the National Research Development and Innovation Office grant (NKFIH; ANN 131305), and the Hungarian Academy of Sciences [Magyar Tudományos Akadémia (MTA)] through the János Bolyai grant (BO/00237/19/2).

## Author contributions

K.I. and R.Baum. conceived the study. K.I. performed the EEG processing and connectivity analyses. R.Baru. performed the statistical analyses and offered insights during the study design. G.D. contributed to concept refinement and offered consultation throughout the implementations. K.I., R.Baru, R.Baum and B.T. designed the data presentation and wrote the manuscript. All authors revised the manuscript and approved the current version.

